# Mass-selective and ice-free cryo-EM protein sample preparation via native electrospray ion-beam deposition

**DOI:** 10.1101/2021.10.18.464782

**Authors:** Tim K. Esser, Jan Böhning, Paul Fremdling, Mark T. Agasid, Adam Costin, Kyle Fort, Albert Konijnenberg, Alan Bahm, Alexander Makarov, Carol V. Robinson, Justin L. P. Benesch, Lindsay Baker, Tanmay A.M. Bharat, Joseph Gault, Stephan Rauschenbach

## Abstract

Electron cryomicroscopy (cryo-EM) and single-particle analysis (SPA) have revolutionized structure determination of homogeneous proteins. However, obtaining high-resolution structures from heterogeneous samples remains a major challenge, as the various protein states embedded in thin films of vitreous ice may be classified incorrectly, resulting in detrimental averaging of features. Here we present native electrospray ion-beam deposition (native ES-IBD) for the preparation of extremely high-purity cryo-EM samples, based on mass selection in vacuum. Folded protein ions are generated by native electrospray ionization, mass-filtered, and gently deposited on cryo-EM grids, and subsequently frozen in liquid nitrogen. We demonstrate homogeneous coverage of ice-free cryo-EM grids with mass-selected proteins and protein assemblies. SPA reveals that they remain structurally intact, but variations in secondary and tertiary structure are currently limiting information in 2D classes and 3D EM density maps. Our results show the potential of native ES-IBD to increase the scope and throughput of cryo-EM structure determination.

## 1 Introduction

Interactions between proteins and other biomolecules govern the cellular processes of life. The most important factors dictating how biomolecules interact with each other are their chemical composition and three-dimensional (3D) structure. Biomolecules perform a wide range of biological functions, reflected by high conformational, and chemical heterogeneity of states. A complete understanding of their function requires selection and visualization of each of these states individually at high resolution.

Most structures in the protein data bank (PDB) were obtained by X-ray crystallography or nuclear magnetic resonance (NMR). X-ray crystallography rapidly produces high-resolution structures from crystalline samples. ^1^ NMR resolves dynamical aspects of structure of smaller biomolecules in their native environment.^2^ Electron cryomicroscopy or cryo-electron microscopy (cryo-EM) allows for high resolution imaging, even if molecules are intractable to crystallization or too large for NMR. However, sample preparation for all three methods is currently limited by solution-based purification. Gas-phase purification using mass spectrometry (MS) and ion-mobility spectrometry (IMS) which can separate molecules by mass to charge at high resolution or collisional cross section provides an extremely powerful orthogonal approach.

TEM protein sample preparation based on native MS and ion-beam deposition (IBD) has been suggested previously.^3^ In pioneering experiments Mikhailov et al. used a modified quadrupole time-of-flight mass spectrometer (Q-ToF MS) and negative stain EM to demonstrate that GroEL and apoferritin remained globular after in-vacuum deposition. However, the sample quality was limited by low ion-beam intensity (below 1 pA), lack of landing energy control (tens of eV per charge), and the use of staining. A systematic characterization of the effects of substrate, landing energy, storage time in vacuum and at ambient conditions by 2D and 3D classification of thousands of particles, now routine in cryo-EM, was not feasible at the time.

In this work, we present a fundamentally new cryo-EM sample preparation workflow, which combines state-of-the-art native MS and electrospray ion-beam deposition (ES-IBD). Native electrospray ion-beam deposition (native ES-IBD) replaces solution-based enrichment and purification with continuous accumulation of mass-filtered biomolecules on cryo-EM grids in vacuum, under controlled deposition conditions. To achieve this, we extended and modified a commercial MS platform (Thermo Scientific^™^ Q Exactive^™^ UHMR) by integrating custom ion-beam deposition hardware and software. With this workflow we were able to prepare ice-free cryo-EM samples of mass-selected, adsorbed, native, gas-phase proteins and protein assemblies. Protein shapes and complex/assembly topologies obtained after imaging and 3D reconstruction were consistent with those obtained from vitrified samples.

Recently, cryo-EM has evolved into one of the leading methods for structural characterization of folded proteins and protein complexes at atomic resolution. ^4,5^ The low contrast of organic matter and its sensitivity to radiation damage and dehydration prevents the use of conventional transmission electron microscopy (TEM) for direct imaging of individual particles at high resolution. While some limitations can be circumvented by negative stain EM, where molecules are embedded in salts containing heavy atoms, this method sacrifices access to high resolution and information on internal protein structure. In a conventional cryo-EM workflow, enriched and purified biological macromolecule solutions are applied to TEM grids, which are then blotted and rapidly submerged into liquid ethane, forming thin layers of vitreous (amorphous) ice and preserving native structure.^6^ Cryogenic temperatures enable access to high-resolution information by immobilizing the sample, preventing dehydration, and retarding the spread of radiation damage.^7^ In single-particle analysis (SPA), high-resolution 3D EM density maps are reconstructed from low-dose, two-dimensional (2D) projection images of thousands to hundreds of thousands of single particles.^8^ Recent technological advances in direct electron detectors, bright electron point sources, energy filters, and aberration-corrected lenses enable for atomic resolution close to 1 Å.^8–13^

However, reliable preparation of chemically pure and homogeneous cryo-EM samples remains one of the most important challenges in cryo-EM, making determination of high-resolution structures challenging and sometimes impossible.^14–20^ Fragile biomolecules can denature during solution-based enrichment and purification, at the substrate-solvent and air-solvent interface, due to blotting, evaporation-induced buffer modification, and during plunge freezing.^14,18,21,22^ As a result, unambiguous classification of single-particle images of to different oligomers, conformers, fragments, and contaminants is not always possible, limiting resolution and hiding conformational flexibility.

Consequently, several methods have been developed to improve cryo-EM sample preparation. In favorable cases, image classification algorithms, can be used to select more homogeneous particle subsets after imaging to obtain multiple structures from a single grid.^23^ Sample heterogeneity can be reduced by crosslinking^24^, introduction of stabilizing ligands,^25^ and sampling of buffers^26^. Functionalized substrates or detergents can be used to keep particles away from the air-water or substrate-water interface. ^27–29^ However, successful application of these approaches is not guaranteed for any given biomolecule.

The native ES-IBD workflow introduced here, is changing sample preparation fundamentally by combining native MS and ES-IBD. Native MS allows to retain proteins in a near-native, folded state in an ultra-pure gas-phase ion-beam, providing information on mass, composition, conformation, and ligand binding sites. ^30^ To this end, intact, non-covalently bound protein complexes are transferred from solution into a mass spectrometer using electrospray ionization (ESI) sources with low flow rates, aqueous solvents, and volatile buffers. Because gas-phase protein ions are separated according to their mass-to-charge ratio (m/z), even proteins from within heterogeneous environments can be examined. For instance, folded membrane proteins or protein complexes, even with entire segments of the cell membrane, can be transferred into vacuum, allowing to purify them or their fragments within the mass spectrometer. ^31^ In addition, native MS enables structurally sensitive techniques like IMS ^32^, hydrogen-deuterium exchange (HDX)^33^, and various forms of ion activation^34^, which provide indirect information on stoichiometry and conformation. In particular, complementary information from native MS can be crucial to guide interpretation and reconstruction of structures from cryo-EM experiments. ^35–37^

Ion-beam deposition (IBD, also referred to as soft/reactive landing or preparative mass spectrometry), employs mass spectrometric techniques to prepare intense and ultra-pure ion-beams of defined energy for controlled deposition onto surfaces in vacuum. It is used, e.g., for fabrication of films of non-volatile molecules and selective preparation of catalytic surfaces. ^38^ ES-IBD in particular has been successfully combined with imaging techniques including scanning probe microscopy (SPM), TEM, and low energy electron holography (LEEH).^3,39–45^ These experiments show that individual peptides and proteins can be separated from solvent and contaminants, deposited, and imaged. Control over ion-beam composition by mass selection and over landing energy allows to address fundamental questions related to gas-phase structures, mechanical properties, and substrate interactions. In this work, ES-IBD forms a direct connection between the selectivity of native MS and the imaging power of cryo-EM, demonstrated by 3D reconstruction of protein assemblies after gas-phase purification and gentle surface deposition.

## 2 Results

We have implemented a native ES-IBD instrument based on a commercial, high-resolution, tandem mass spectrometer, designed for analysis in a mass range up to 80,000 m/z (Thermo Scientific Q Exactive UHMR). We modified the ion-source, instrument operation mode, and added home-made deposition ion-optics and control software (see Fig. S1). The modifications allow for extraction of an ion beam from the collision cell for controlled deposition onto various substrates, including TEM grids, at room temperature and in vacuum. We demonstrate homogeneous coverage of ice-free cryo-EM grids with mass-selected specimens. Ion-beam intensities for mass-selected native proteins range from 10 to 100 pA, and typically allow cryo-EM grids to be covered within 30 minutes. Careful thermalization and guiding of the ion-beam allows to keep landing energies below 2 eV per charge, where covalent bonds are unlikely to be affected.^38,46,47^ The significant improvements of ion-beam intensity and landing energy control, combined with freezing in liquid nitrogen after sample transfer under ambient conditions, and imaging at cryogenic temperatures, enabled us to demonstrate 3D reconstruction after gas-phase purification and gentle surface deposition.

### A native ES-IBD workflow for structure determination

Fig. 1 gives an overview of the complete sample preparation workflow from solution to 3D EM density map (see Methods for details on used materials, protein preparation, native MS, and SPA). Protein solutions are prepared using a standard native MS workflow, including exchange of buffer to volatile ammonium acetate. ^48^ To achieve the highest m/z precision, ions are usually desolvated using collision induced dissociation (CID) in the atmosphere-to-vacuum interface or higher-energy collisional dissociation in the quadrupole collision cell. Instead, for all experiments presented here, we use reduced electric potential gradients throughout the instrument to promote soft, non-activating conditions, while retaining high ion transmission.

**Figure 1:**
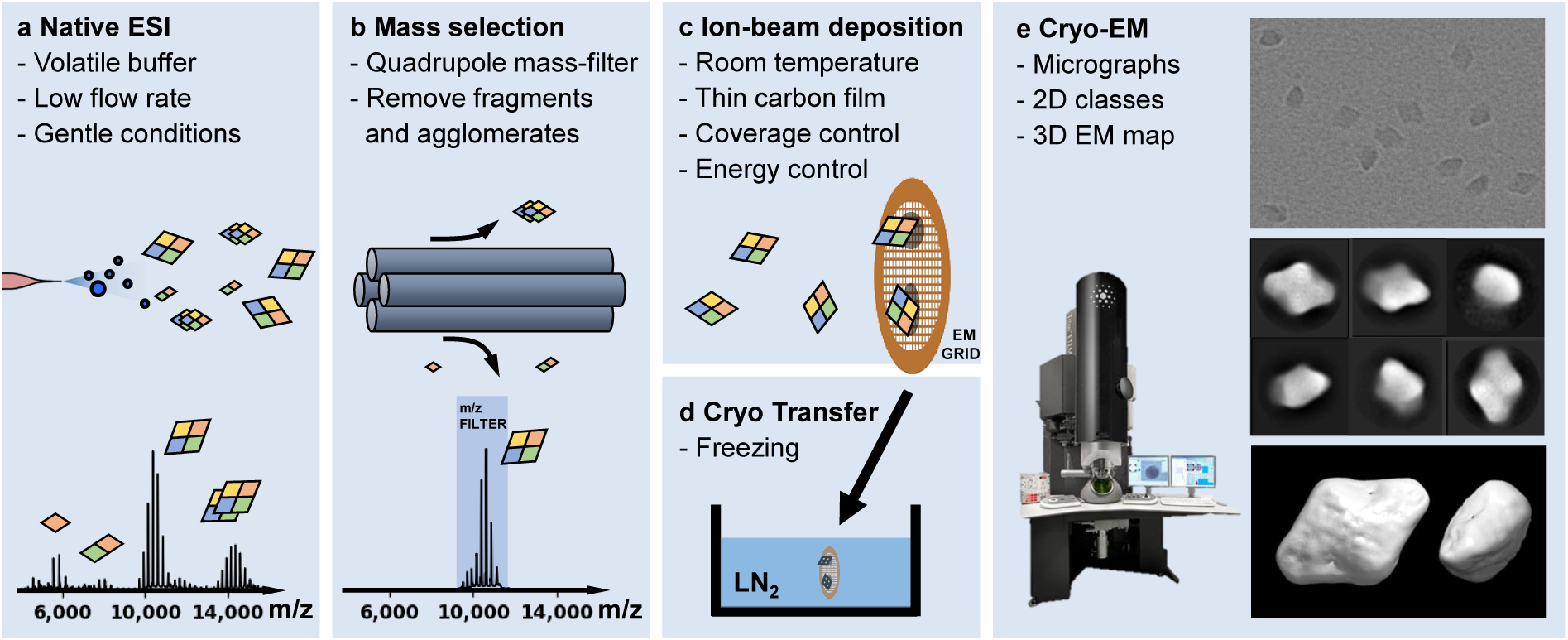
Schematic representation of mass-selective cryo-EM sample preparation using the native ES-IBD workflow. Mass spectra, micrographs, 2D classes, and 3D EM density maps shown here were obtained by applying the workflow to *β*-galactosidase (compare Fig. 4). **a** Proteins are transferred into the gas phase via native electrospray ionization. **b** The species of interest is mass-selected and separated from fragments, aggregates, and contaminants. **c** The mass-selected ion-beam is deposited with defined landing energy on TEM grids. **d** Grids are removed from deposition chamber and plunged in liquid nitrogen, followed by cryogenic transfer to the EM. **e** Samples are imaged and micrographs are processed according to established SPA procedures, resulting in 2D classes and 3D EM density maps.

Initially, mass spectra are acquired to ensure native ionization conditions, evaluate nano-electrospray stability, and to select a suitable quadrupole mass filter transmission range. The instrument mode is then switched, to guide the ion beam through the collision cell, where it is thermalized, before entering a custom deposition stage and reaching a custom, room-temperature sample holder at a pressure of 10^−5^ mbar. An integrated, retarding-grid ion-current detector records the ion-beam intensity, total beam energy, and beam energy distribution. The concentration of the protein solution and ion-source parameters are optimized for beam intensity and stability. The total beam energy (kinetic energy and potential energy relative to ground) typically decreases by only 1.5 eV per charge between the collision cell and sample, due to collisions with the background gas, indicating non-activating transfer through the custom DC ion optics. The full-width-half-maximum (FWHM) of the beam energy distribution is typically below 1 eV per charge, allowing to use landing energies below 2 eV per charge.

Prior to deposition, TEM grids with carbon films are placed into the sample holder, and inserted into the deposition chamber via a vacuum load-lock. Plasma-cleaning to produce a more hydrophilic surface is not required, as our method bypasses the application of a liquid to the grid at any point. The deposition current on the grids is monitored using a picoammeter to align the ion beam and obtain the total deposited charge, in picoampere hours (pAh), by integrating the deposition current. Typically, 10 to 15 pAh are collected in 30 minutes. The total deposited charge and mean charge state are used to estimate the number of deposited proteins, typically about 10^10^ particles, and achieve consistent coverage between experiments. Particle density measurements across the grid typically indicate a Gaussian-shaped particle density distribution with a FWHM around 1 mm, meaning that the entire grid is covered without moving the beam and the particle density increases towards the grid center.

After deposition, TEM grids are retrieved via the vacuum load-lock, transferred under ambient conditions, and manually plunged into liquid nitrogen within two minutes. Note that fast cooling rates as used for vitrification are not required due to the absence of water. Samples are stored in liquid nitrogen for less than two days. Subsequent sample transfer, imaging, and data processing proceeds using standard SPA workflows established for samples with vitrified ice.

### Preparation of clean and homogeneous samples of high density

Initial experiments were conducted with deposition and imaging at room-temperature, i.e. without freezing. We deposited an apo/holo-ferritin mixture on Quantifoil® grids with homemade graphene oxide (GO) films. The high contrast of the iron core of holoferritin allowed investigation of particle density and distribution at room temperature, even though the protein was hardly visible. Fig. 2a shows a variation of particle density with GO substrate film thickness across a single micrograph. The particle density is high and homogeneous on thick GO films (left), but only few particles are found on thin films (right). We rationalize this observation by suppression of thermal particle diffusion due to stronger Van-der-Waals interaction and higher density of edges and defect sites on the thicker areas, as previously observed for the adsorption of clusters on freestanding graphene. ^49^

**Figure 2:**
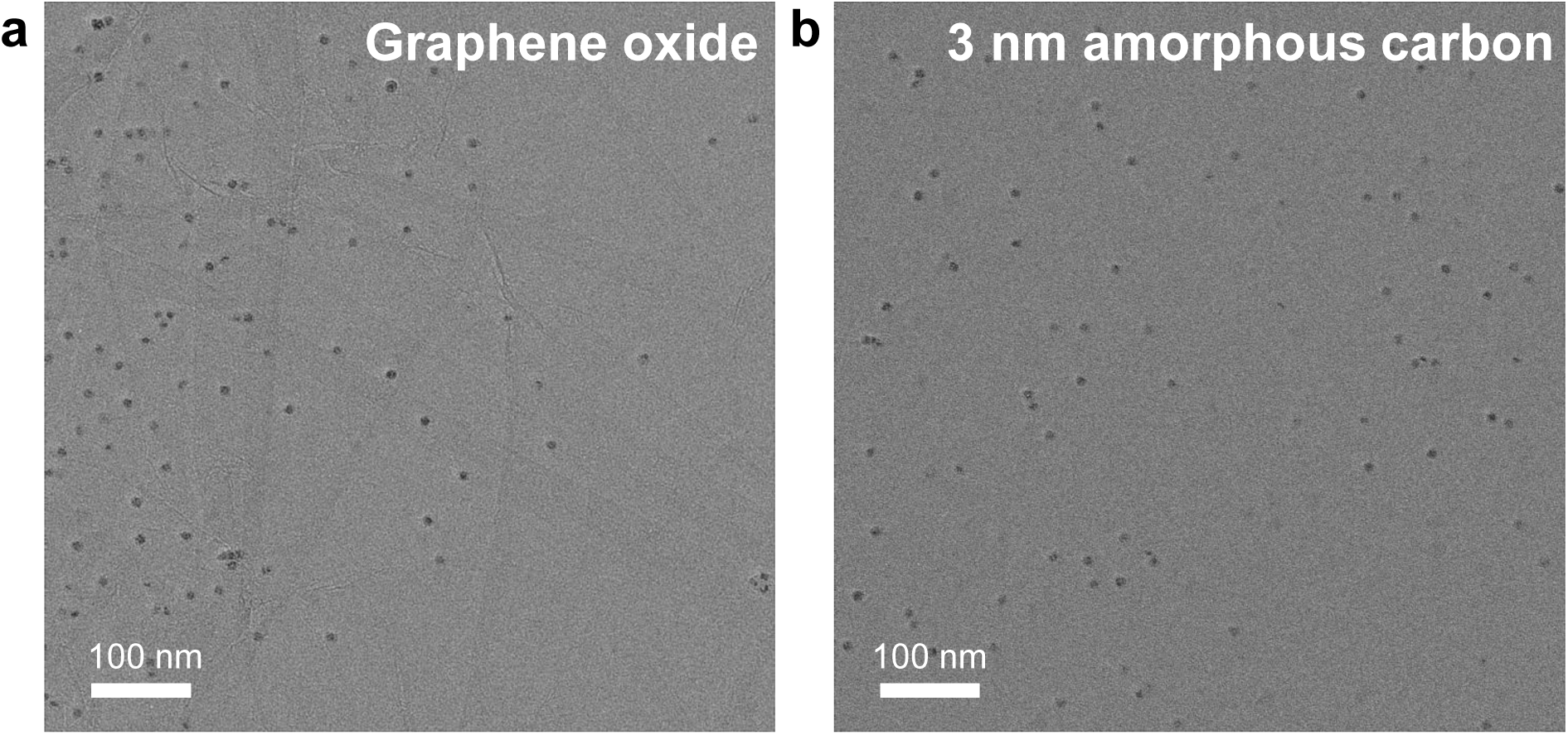
**a** Room-temperature TEM micrograph of ferritin, deposited on a home-made graphene oxide film. Particles are found mainly in areas with thick GO film (left side). **b** Room-temperature TEM micrograph of ferritin on a 3 nm amorphous carbon film under identical deposition and imaging conditions, demonstrating a clean sample with homogeneous particle density.)

To avoid artifacts caused by the irregular surface and film thickness of graphene oxide films, we proceeded to use commercial 3 nm amorphous carbon grids which suppress thermal diffusion and provide a consistent homogeneous background in EM micrographs over the entire grid. This enabled us to obtain homogeneous coverage as shown in Fig. 2b. The majority of particles are well separated from each other rather than aggregated. The entire grid is covered with increasing particle density towards the center, allowing to select an ample number of grid squares with ideal density for data collection.

### Protein structure and subunit topology is retained after gas-phase purification and deposition

We then added a manual freezing step in liquid nitrogen directly after deposition and transfer under ambient conditions. Cryogenic temperatures were then maintained throughout the transfer to the microscope and imaging. We applied this workflow to prepare and image native ES-IBD samples for a range of protein assemblies, including apo/holo-ferritin (24-mer), alcohol dehydrogenase (ADH, tetramer), *β*-galactosidase (*β*-gal, tetramer), and GroEL (tetradecamer). These molecules cover a mass range from 147 kDa (ADH) to 803 kDa (GroEL), have characteristic shapes, and corresponding high-resolution cryo-EM density maps are available in the PDB. For each sample, we collected 50 micrographs, resulting in up to 3,000 single particles. We performed 2D classification in RELION 3.1 (see Methods for more details). The resulting micrographs are shown in Fig. 3, along with 2D classes and representative single particles as well as corresponding 3D models from the PDB and published 2D classes ^13,50,51^, obtained from traditional plunge-frozen cryo-EM samples of protein in vitrified ice from grids with holey carbon films.

**Figure 3:**
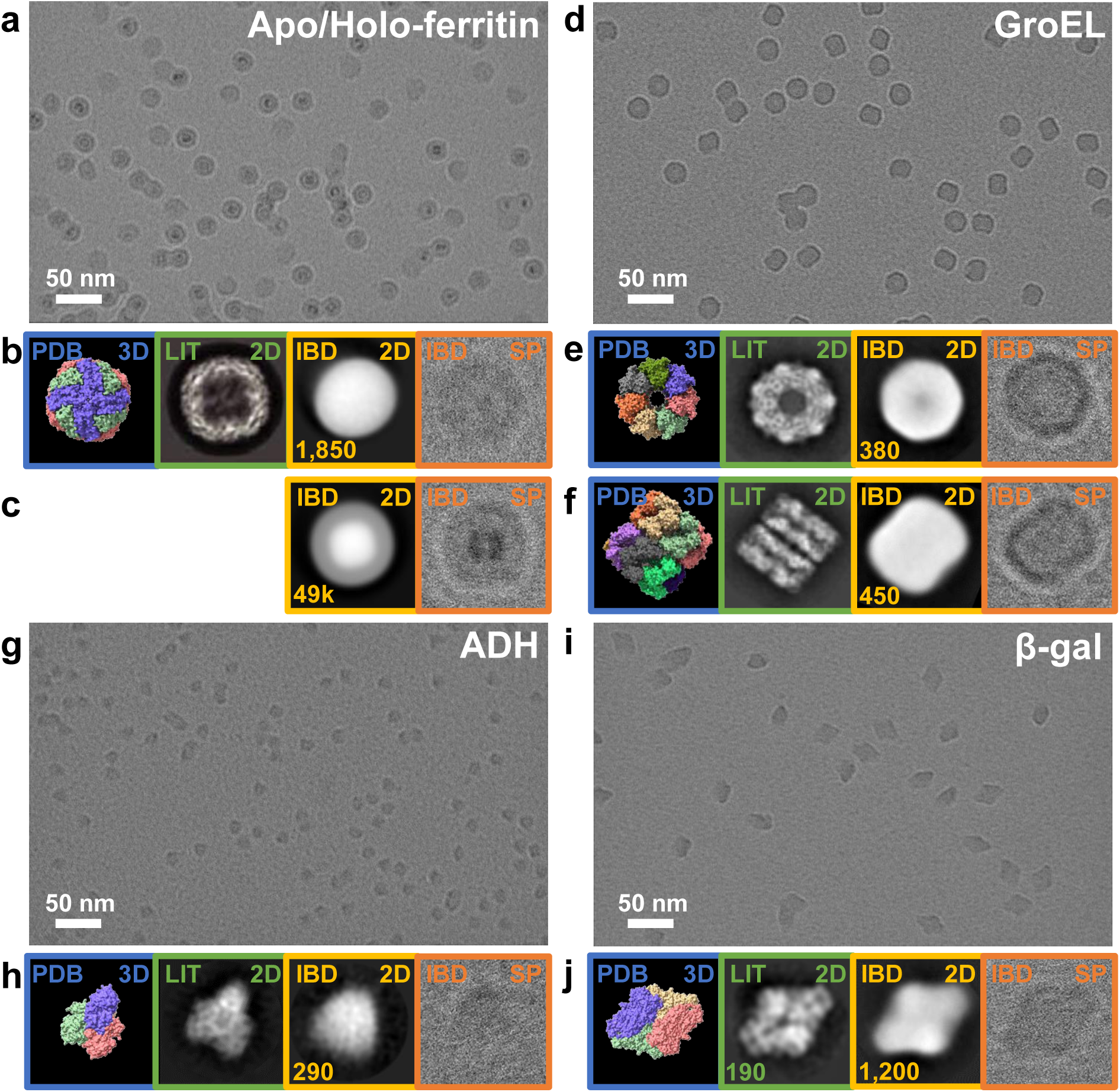
Unfiltered cryo-EM micrographs of ice-free native ES-IBD samples: **a** apo/holo-ferritin, **d** GroEL, **g** ADH, and **i** *β*-gal. Panels beneath the micrographs show, from left to right, 3D models from the PDB (blue, PDB 3D), 2D classes from plunge-frozen cryo-EM samples from the literature (green, LIT 2D), and 2D classes (yellow, IBD 2D) and representative single particles (orange, IBD SP) form ice-free native ES-IBD samples. Panels **b** and **c** show apo- and holo-ferritin, panels **e** and **f** show GroEL top and side views, and panels **h** and **j** show ADH and *β*-gal. 3D PDB models were rendered with ChimeraX^54^ using PDB entries 7A6A (apoferritin), 5W0S (GroEL), 7KCQ (ADH), and 6CVM (*β*-gal). 2D classes from literature were taken from Yip et al. (apoferritin), Roh et al. (GroEL), Guntupalli et al. (ADH), and the RELION 3.0 tutorial data set (*β*-gal). The number of particles in the 2D classes is11given where available.

Fig. 3a shows a sample prepared by depositing an apo/holo-ferritin mixture. In contrast to the room temperature micrographs, the ferritin protein shells are clearly visible around the iron cores when imaging samples at cryogenic temperatures after deposition. Apoferritin, barely visible when imaging at room temperature, can be clearly identified as well. The single apoferritin particle in Fig. 3b appears circular, but the corresponding 2D class has a slightly oval shape, suggesting a deformation of the protein shell during the workflow. For holoferritin, Fig. 3c, the single particle shell is round on the outside and rectangular on the inside, closely resembling the PDB structure shown for apoferritin. This is confirmed in the 2D class but further analysis is inhibited as the classification algorithm is confounded by the high-contrast iron core. Interestingly, there are no indications of a localized deformation of the protein shell as for apoferritin. This result is consistent with previous IMS data, suggesting a gas-phase collapse of the apoferritin cavity and stabilization of holoferritin by the iron core. ^52,53^

Fig. 3d shows a micrograph of a GroEL sample. A very high contrast is observed due to the high mass of 803 kDa and absence of ice. Top and side views can be identified unambiguously. The 2D classes of top and side views clearly show features of the characteristic barrel shape that are already apparent in the single particle images, including the central cavity, a heptameric symmetry in the top view, and the notch between the two heptamer rings in the side view. Further substructure, as present in the literature 2D classes, is not visible.

A micrograph of a sample of ADH, the lowest mass protein complex in our study at 147 kDa, is shown in Fig. 3g. It demonstrates the remarkable contrast of ice-free samples, even for small proteins. Individual particles show characteristic triangular shapes which become much clearer in the 2D class shown in 3h. However, due to a lack of homogeneous internal structure in the single-particle images, specific orientations could not be assigned unambiguously.

Finally, Fig. 3i shows a micrograph of a sample of *β*-gal, which is commonly used as a standard in cryo-EM due to its relatively high mass (466 kDa), high stability, and characteristic shape. In contrast to apo/holo-ferritin and GroEL, *β*-gal has no iron core or cavity. Again, individual particles and their orientations can be identified unambiguously from the unfiltered micrograph. The most prominent diamond-shaped 2D class and a corresponding single particle are shown in Fig. 3j.

Generally, there is no evidence for unfolding into extended polypeptide chains or fragmentation into subunits. The significantly improved contrast and well-defined shape in the 2D classes of the ice-free samples indicate consistency of low-resolution structural features among particles and preservation of quaternary structure. Particle dimensions in 2D classes indicate no lateral deviation from literature values. However, there is a consistent lack of defined internal structure compared to the 2D classes from conventionally prepared samples with vitreous ice.

### 3D reconstruction based on native ES-IBD samples

To assess the currently accessible structural information in more detail, we collected approx. 3,000 movies of *β*-gal from which we obtained 50,000 single particles images, a significantly larger data set than above. For direct comparison, we prepared a control sample by applying the solution used for native ES-IBD directly to a grid with holey carbon film, followed by conventional blotting and plunge freezing. Approx. 900 movies were recorded with identical microscope settings (see Methods), resulting in 93,000 particles.

Fig. 4 shows unfiltered EM micrographs, band-pass filtered EM micrographs, 2D classes, and 3D EM density maps of both samples. As described above, the particle contrast for the native ES-IBD sample is very high, the particle density is very homogeneous, and individual particles are well separated from each other. In contrast, the unfiltered micrograph of the vitrified control sample shows agglomerated particles. Individual particles and orientations are hard to identify by eye and only become visible after band-pass filtering. After 2D classification, randomly distributed particle orientations, shown in Fig. 4f, are obtained for the control sample, whereas strong preferred orientation is observed for the native ES-IBD sample. The latter exclusively shows three orientations, shown in Fig. 4e, with a particle percentage of 56 % (diamond-shaped), 43 % (tilted), and *<* 1 % (top).

**Figure 4:**
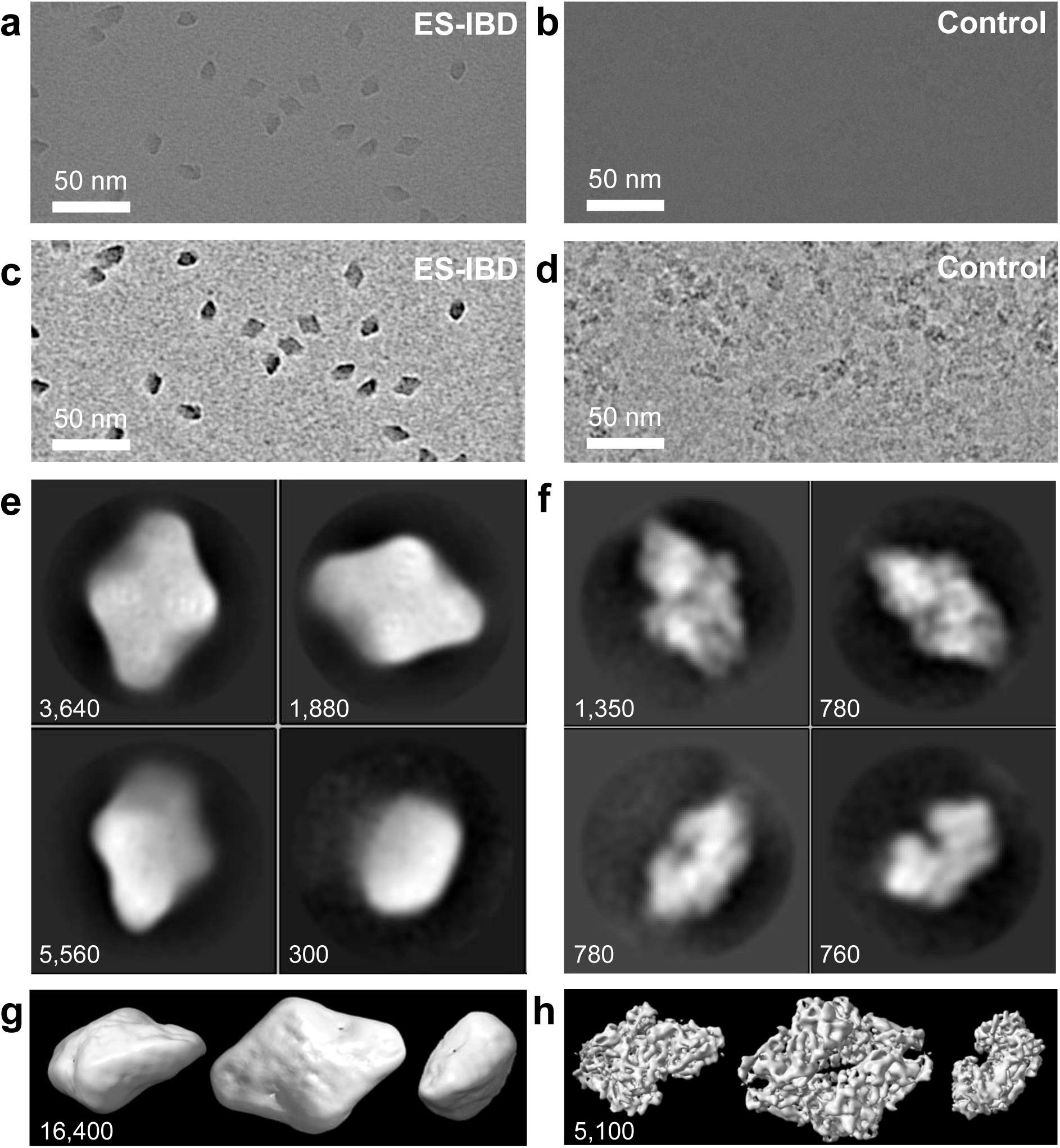
Unfiltered EM micrographs, **a** and **b**, EM micrographs band-pass filtered between 10 and 200 Å, **c** and **d**, representative 2D classes, **e** and **f**, and 3D EM density maps, **g** and **h**, for a native ES-IBD and a plunge frozen control sample of *β*-gal, respectively. The number of particles in the 2D classes and 3D maps is indicated in the panels.

2D classes of the control sample show features of strong internal contrast, corresponding to secondary structure. These features are missing in the native ES-IBD sample, despite a much better contrast relative to the background. However, a characteristic symmetric pattern can be observed in the diamond-shaped 2D classes. It consists of clusters of 4 high-density spots, four low-density spots around the center as well as at both tips. The smallest distance between the high-density spots is 8 Å. In addition, two of the opposing diamond edges have lower contrast than the other two (see Fig. S2 in the SI for marked features).

Next, we look at the 3D EM density maps of the ES-IBD and control sample in Fig. 4g and h, respectively. The level of information is consistent with that in the 2D classes. For the control sample, a 9 Å (0.143 Gold-standard FSC) 3D EM density map was obtained and shows very good agreement with the secondary structure from the PDB (see Fig. S4). The corresponding 3D map from the native ES-IBD sample reliably reproduces the external features but lacks internal detail (see Fig. S5 and Movie 1 in the SI). To validate the results from SPA, we have also performed electron cryotomography (Cryo-ET) on a separate native ES-IBD sample to which gold fiducials were added before deposition. The resulting tomogram is in good agreement with the 3D classes from SPA (see Fig. S6 and Movie 2 in the SI). It shows the same well-defined external and less defined internal features.

## 3 Discussion

Our results clearly show that proteins remain folded and subunits in protein complexes remain attached to each other in their native orientation during sample preparation using native ES-IBD, despite dehydration, collision with the substrate, and prolonged exposure to the substrate-vacuum interface at room temperature. However, the resulting sample heterogeneity is currently limiting access to secondary structure. For *β*-gal, sub- nanometer resolution is routinely achieved using conventional vitrified samples, but this is not currently attained with native ES-IBD samples. In the following we discuss the main differences between the two sample preparation methods that can explain this discrepancy.

First, the successful generation of a subnanometer 3D map for the control sample shows that neither the preparation of the solution for mass spectrometry, the microscope performance and used settings, nor the number of collected single particle images is limiting access to secondary structure.

Second, the mass spectrum indicates that less than 100 water molecules remained attached to the gas-phase protein ion (based on spectral deconvolution and mass measurement), i.e., only a small fraction of the 4,194 structural water molecules in the PDB model. MD simulations and gas-phase experiments indicate that dehydration can lead to a range of kinetically trapped gas phase structures, however, the native secondary structure is typically largely preserved and gas-phase proteins are able to refold at least partially upon rehydration.^55–57^ In particular, it has been shown that enzymes can retain their activity after deposition onto liquid surfaces. ^58^

Third, particles may be damaged by the collision with the carbon film. In the present case, *β*-gal molecules land with a kinetic energy of 2 eV per charge, corresponding to a velocity of about 200 m/s. The nature and severity of damage caused by this type of collision are currently poorly characterized. Previous studies show that covalent bonds are unlikely to be affected or broken,^38,46,47^ but conformation can change. ^59^ Localized damage of the protein surface and random orientations on impact could introduce sample heterogeneity that leads to loss of information during averaging. It is intuitive to expect a larger deformation when a protein lands on a sharp edge rather than a flat plane. Indeed, when allowing for asymmetry in the 3D reconstruction for the ice-free native ES-IBD sample, we obtained classes that show a localized indentation close to one of the narrow tips of *β*-gal (see Fig. S3 in the SI).

Fourth, the protein samples spend up to 30 minutes at the substrate-vacuum interface at room-temperature before being frozen. This highly asymmetric environment can cause additional, thermally mediated deformation. E.g., the low-contrast edges in the diamond-shaped class of *β*-gal (see Fig. S2c) possibly indicates a stronger deformation of the two subunits that are in direct contact with the substrate. The effect of the substrate-vacuum interface on particle orientations and structural integrity has been characterized for a broad range of materials.^38^ Its effect on proteins however, is still poorly understood and challenging to control. Further work is required to characterize the effects of various substrates and surface modifications on thermal motion, sample stability, and preferred orientation.

Fifth, particles can rehydrate during the brief transfer at ambient conditions in our current workflow. The limited control of the degree of rehydration in this step is a major limitation that needs to be addressed in the future.

Sixth, while the dependence of radiation damage on temperature has been investigated thoroughly,^60^ there is little research on the role of the ice layer, as direct comparison with ice-free samples was previously not possible. Likewise, the degree of damage due to dehydration has not been investigated systematically. Our workflow allows to address these effects directly, to gain a better understanding of the role that molecular waters play in these processes.

Finally, it is worth considering that data processing in the SPA workflow is highly optimized for vitrified samples and may require adjustment to obtain more information from ice-free native ES-IBD samples.

We conclude that in our workflow, the overall shape of protein assemblies and with it the secondary structure is largely preserved, but we introduce heterogeneity in the secondary and tertiary structure, by dehydration, landing, and surface interactions which limits the amount of information that can be obtained by averaging techniques. A thorough characterization of the individual aspects discussed above is needed to improve sample quality and thus ultimately achievable experimental resolution.

## 4 Conclusions

We have implemented native electrospray ion-beam deposition (native ES-IBD) as a novel approach for reproducible preparation of ice-free, homogeneously covered, and high-density cryo-EM samples. This method provides sample purity by mass-selecting a native protein ion-beam in a mass spectrometer. Protein conformation is largely preserved during transfer onto a TEM grid by carefully controlled native MS and soft landing. With our prototype instrument we demonstrated the fabrication of cryo-EM samples for a diverse selection of protein assemblies (apo/holo-ferritin, GroEL, ADH, and *β*-gal). Particles stand out clearly against the background, even for proteins with a molecular weight around 150 kDa, and different orientations can typically be distinguished by eye in unfiltered micrographs. Further, we showed that 3D reconstruction from ice-free samples is possible using SPA.

The extensive optimization of the solution-based purification and vitrification is replaced by optimization of native MS conditions, which typically requires less expensive instrumentation, provides faster feedback, and offers orders of magnitude higher selectivity. Due to the modular design of our deposition stage, the workflow can be modified to allow preparation of conformation-selective samples using IMS. Significant improvements in ion transmission are possible^61^ and promise fast preparation of ultra-pure samples of individual complexes, charge states, and even ligand bound states. Collection of sufficient particles for image reconstruction on a chromatographic timescale also appears possible. As we based our workflow on a commercial instrument, all these abilities can be transferred to other labs with reasonable effort.

Even with its currently limited resolution, the mass-spectrometric information, selectivity, and strong contrast even in the unfiltered micrographs can be useful for screening and interpretation of higher resolution structures obtained using conventional cryo-EM. Further, native ES-IBD can provide complementary information to help address the challenges of the plunge freezing workflow discussed in the introduction. There may be great potential in eliminating solvent and ice-related issues, including denaturation at the air-solvent interface, strong and inhomogeneous background signal, unintentional devitrification, beam-induced motion of the ice, and inhomogeneous particle distribution.

Imaging proteins after controlled gas-phase exposure allows to address fundamental questions on the nature of the native-like gas-phase state, widely debated in native MS and IMS.^53,62,63^ The interaction of a protein with different solid surfaces in a hyperthermal collision is of great interest, as it provides a direct means to probe mechanical properties of the protein. While we typically perform experiments in the soft-landing regime to maintain biochemically relevant stoichiometry and native conformation, activation can be readily achieved, either within the mass spectrometer or by reactive landing or surface induced dissociation, to image collision-energy specific states and fragments ^47,59^.

After 40 years in which the underlying principles of cryo-EM specimen preparation remained essentially unchanged,^16,20^ native ES-IBD enables a variety of truly novel workflows. Establishing cryo-EM with samples from mass-selected protein deposition provides a great variety of research opportunities for structural biology, because it directly relates chemical information to biological structure.

## 5 Methods

### Protein preparation

Soluble proteins, alcohol dehydrogenase (ADH, A7011-15KU), *β*-gal (*β*-gal, G3153-5MG), ferritin (F4503-25MG), and GroEL (chaperonin 60, C7688-1MG), were purchased from Sigma Aldrich and used without further purification unless otherwise specified. Ammonium acetate (7.5 M, A2706-100ML), MeOH (1060352500), Acetone (1000145000) and buffer components for the reconstitution of GroEL, Tris (93362-250G), KCl (P9541-500G), EDTA (DS-100G), MgCl2 (63068-250G), ATP (A6419-1G), were also purchased from Sigma Aldrich. All concentrations are calculated with respect to the most abundant oligomers. Lyophilized powders of alcohol dehydrogenase, *β*-gal, and were resuspended in 200 mM ammonium acetate (pH 6.9) to a final concentration of 50 *µ*M. The saline ferritin stock solution had a concentration of 260 *µ*M.

GroEL was reconstituted from lyophilized powder. To this end, the powder was dissolved in 100 uL MeOH and 400 uL buffer, containing 20 mM Tris, 50 mM KCl, 0.5 mM EDTA, 5 mM MgCl2, 0.5 mg/mL ATP, mixed for 2 hours at room temperature, precipitated by adding 500 uL cold Acetone and centrifuged to form a pellet. After disposing of the supernatant, the pellet was resuspended in 250 uL of the original buffer and gently mixed overnight. The final solution had a concentration of 5 *µ*M and was aliquoted and stored at -80 °C until use.

All proteins were desalted by passing through two P6 buffer exchange columns (7326222, Biorad), equilibrated with 200 mM ammonium acetate (pH 6.9). If applicable, they were then diluted in 200 mM ammonium acetate (pH 6.9) to reach the concentration used for native MS: 5 *µ*M (alcohol dehydrogenase), 10 *µ*M (*β*-gal), 8 *µ*M (ferritin) 5 *µ*M (GroEL). Buffer exchange was always done on the day of deposition.

### Native mass spectrometry

Borosilicate glass capillaries (30-0042, Harvard Bioscience) were pulled into nano electrospray ionization emitters, using a pipette puller (P-1000, Sutter Instrument), and gold coated using a sputter coater (108A/SE, Cressington). For native MS, up to 10 *µ*L of protein solutions was loaded into an emitter and the emitter tip was clipped to produce an opening of 0.1 to 10 *µ*M.^64^ Electrospray ionization was initiated by applying a potential of 1.2 kV and a gentle nanoflow pressure (*<* 200 mbar above atm). Modifications of the ion source are described in Fig. S1.

General instrument conditions were as follows: Source DC offset = 21 V, S-lens Rf level = 200 V, transfer capillary temperature = 60 °C, ion transmission settings set to ”High m/z” (700 Vp-p for the injection flatapole, and 900 Vp-p, for the bent flatapole, transfer multipole, and collision cell), detector optimization ”High m/z”, injection flatapole = 5 V, interflatapole lens = 4 V, bent flatapole = 2 V, transfer multipole = 0 V, collision-cell pressure setting 7 (N_2_), collision-cell multipole DC -5 V, collision-cell exit-lens -15 V. For collection of mass spectra the instrument was operated in standard mode, ^65^ and for ion deposition, a modified scan matrix was used that allowed ions to pass through the C-trap and collision cell directly into the deposition stage, without trapping. Mass spectra are shown in Fig. S7.

### TEM grids

Cu TEM grids with mesh size 400 were purchased from Agar Scientific, including 3 nm amorphous carbon on a lacey carbon film (AGS187-4) and holey carbon film (AGS174-3). A graphene oxide layer was added to the latter by plasma-cleaning for five minutes, drop casting of 3 *µ*l graphene oxide suspension (763705-25ML, Sigma Aldrich), diluted in water to 0.2 mg/ml, blotting with filter paper (11547873, Fisherbrand) after one minute, followed by three washing and blotting steps with water. No further treatment was applied to grids before deposition.

### Plunge-freezing control sample

The control sample for *β*-gal was prepared using ammonium acetate solutions used for native ES-IBD and a Cu/Rh 200 mesh grid (Q2100CR2, Quantifoil®) with 2 *µ*m holes and 2 *µ*m spacing between the holes. 3 *µ*L of a 5 *µ*M solution was applied to the grid, followed by blotting and plunging into liquid ethane, using a Vitrobot (Thermo Fisher Scientific) at a relative humidity of 100 % and a temperature of 10 °C.

### Image acquisition and processing

All micrographs were collected using microscopes at the COSMIC Cryo-EM facility at South Parks Road, University of Oxford, UK. Room temperature screening was done on a Talos F200C (Thermo Fisher Scientific) cryo-TEM with Ceta detector. Micrographs of native ES-IBD samples of apo/holo-ferritin as well as tomographic tilt series of *β*-gal were imaged on a Titan Krios 300 kV cryo-TEM (Thermo Fisher Scientific) with a BioQuantum energy filter and a K3 direct electron detector (both Gatan). All other data was acquired on a Talos Arctica 200 kV cryo-TEM with a Falcon 4 direct electron detector (both Thermo Fisher Scientific). Manual and automated data acquisition were controlled using EPU software (Thermo Fisher Scientific).

All micrographs were recorded using a range of defocus settings between -1 and -3 *µ*m. For the unfiltered micrographs shown in this work, the color range was adjusted to the data range, but no data was cut off and no non-linear adjustments were applied. Typically, 50 micrographs were recorded per sample giving up to 3,000 particles and the magnification was 180E3 corresponding to a pixel size of 0.78 Å.

For *β*-gal, two larger data sets of 3,000 and 900 EER^66^ movies were collected for the native ES-IBD and control sample, providing 50,000 and 90,000 particles, respectively. The magnification for this collection was 240E3 corresponding to a pixel size of 0.59 Å and the total exposure was 40 *e/*Å^2^.

For SPA, data was processed using RELION 3.1.^67^ Motion-corrected MRC files were generated from EER files, using RELION’s own implementation of the MotionCor2 algorithm.^68^ Contrast Transfer Functions (CTFs) were estimated using CTFFIND 4.1^69^ and used for CTF correction in the following steps. The high contrast obtained for native ES-IBD samples allowed for reliable automated particle picking based on a Laplacian-of-Gaussian (LoG) filter. Particles were extracted in 256 × 256 pixel boxes scaled down by a factor of 2. An initial 2D classification step was used to remove incorrectly picked particles and subsequent 2D classification produced the 2D classes shown in the main text.

A *β*-gal model from the PDB (6CVM), low-pass filtered to 60 Å, was used as an initial model for 3D classification. 3D density maps were obtained from the native ES-IBD and control EER data set, using a subset of 16,400 and 5,100 particles, respectively, selected after multiple classifications. Particles were re-extracted in 560 × 560 pixel boxes and downscaled to 256 × 256 pixel boxes. The best-looking 3D class from the last classification was used as reference map, and scaled accordingly using the relion image handler program. The final structures were then produced using RELION’s automated refinement. Videos of the resulting 3D EM density maps were generated using ChimeraX^54^.

### Tomography

A native ES-IBD sample of *β*-gal was prepared using the standard workflow, apart from adding gold fiducials (FG 10 nm, UMC Utrecht) before deposition. To this end 1 *µ*g of gold fiducials was dissolved in 20 *µ*l of water and 2 *µ*l were applied to a plasma-cleaned grid which was then blotted after 30 seconds.

Tilt series were acquired at a target defocus of 8 *µ*m and with a pixel size of 3.47 Å using the SerialEM software^70^ in low-dose mode, using a dose-symmetric tilt scheme. The K3 detector was operated in super-resolution counting mode using dose fractionation. The target tilt range was set to 130° (±65°), with increments of 1°, and a target total electron dose of 130 *e/*Å^2^ was used.

Cryo-ET data analysis was performed in IMOD^70^ using techniques described previously^71^. In brief, movies were aligned and dose-weighted using MotionCor2.^68^ Tilt series alignment was performed using gold fiducial tracking in IMOD^72^. CTF was corrected using ctfphaseflip in IMOD. Tomographic reconstruction was carried out using the SIRT algorithm implemented within Tomo3D^73^, and visualized using ImageJ^74^.

## 6 Acknowledgement

We acknowledge support from Thermo Fisher Scientific who provided the Q Exactive UHMR mass spectrometer within the framework of a technology alliance partnership. A.B, A.M., K.F. and A.K are employees of Thermo Fisher Scientific. T.K.E. acknowledges funding from the European Union’s Horizon 2020 research and innovation programme under the Marie Sklodowska-Curie grant agreement No 883387. T.A.M.B. is a recipient of a Sir Henry Dale Fellowship, jointly funded by the Wellcome Trust and the Royal Society (202231/Z/16/Z). T.A.M.B would like to thank the Vallee Research Foundation, and the Leverhulme Trust for support. J.G. was supported by a Junior Research Fellowship at The Queen’s College, Oxford. This research was funded in whole, or in part, by the Wellcome Trust Grant number 104633/Z/14/Z. For the purpose of open access, the author has applied a CC BY public copyright licence to any Author Accepted Manuscript version arising from this submission. We thank Prof. Susan Lea for fruitful discussions and support with TEM imaging and Dr. Josh Gilbert for valuable feedback on the manuscript.

## Notes

### Competing Interest Statement

The authors have declared no competing interest.

