## SUPPLEMENTARY INFORMATION for "Mass-selective and ice-free cryo-EM protein sample preparation via native electrospray ion-beam deposition"

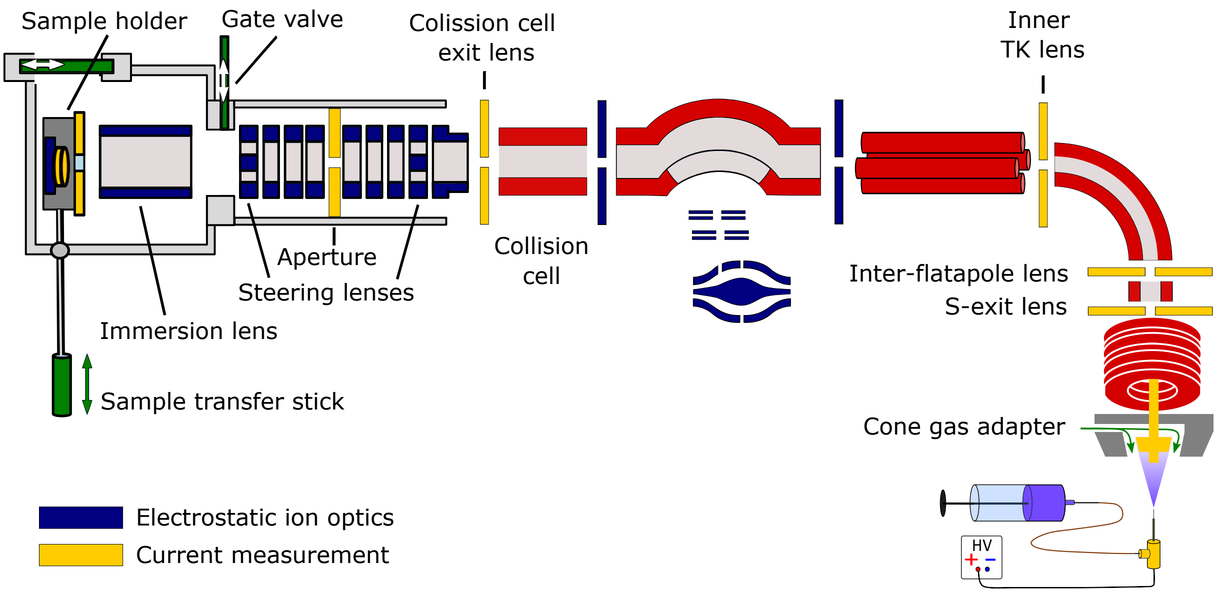


Figure S1: Schematic of the ES-IBD deposition instrument, consisting of a commercial, high-resolution mass spectrometry platform (Thermo Scientific Q Exactive UHMR, right side) and home-made landing stage and sample holder (left side). The instrument combines RF (red) and DC (blue) ion optics. DC ion optics color-coded in yellow are connected to picoammeters (Model 9103, RBD Instruments), to allow for current measurement to optimize ion-beam transmission. The standard transfer capillary with an i.d. of 0.58 mm was exchanged for a custom capillary (based on 590129, CS Chromatographie) with an i.d. of 0.75 mm, which moderately increased transmission. The aperture of the S-exit lens was increased from 1.4 to 2.5 mm, increasing transmission up to tenfold. An additional pump (XDS35, Edwards) was installed as a dedicated backing pump for the source turbomolecular pump to account for the additional gas load. Ions are thermalized in the collision cell at a pressure of approx. $10^{-2}$ mbar. A beam collector (electrometer), downstream of the collision-cell exit-lens, was removed to allow for continuous transmission of the ion beam into the landing stage at a pressure of approx. $10^{-5}$ mbar. The latter allows the beam to be focused and steered onto the sample holder using only DC ion-optics. The sample holder comprises a retarding-grid ion-current detector, which records the ion-beam intensity, total beam energy, and beam energy distribution, as well as two sample positions with variable DC potential, which allows the landing energy to be controlled and to monitor the total charge deposited.

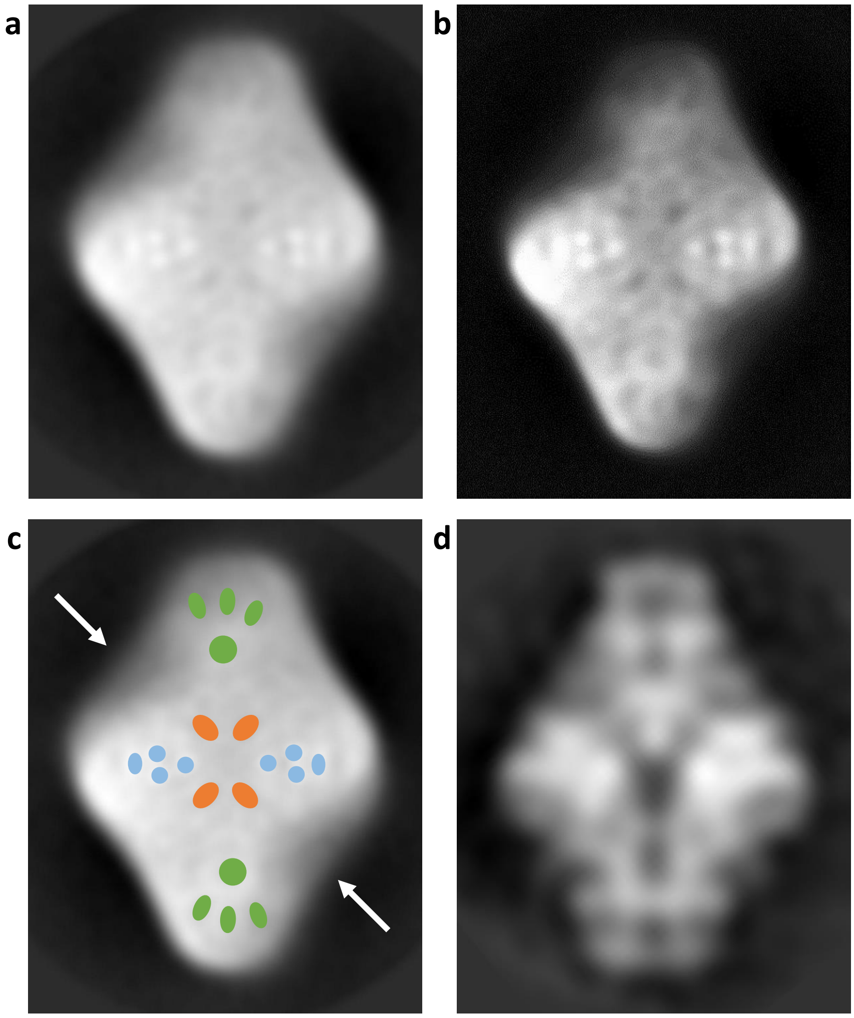


Figure S2: **a** Detailed view of the diamond-shaped class from the ice-free ES-IBD $\beta$-gal sample. **b** Same view with increased contrast and reduced brightness to highlight characteristic internal features. **c** Same view with marked features, including eight characteristic density maxima (blue), four minima around the center (orange), and eight minima at the tips (green). Arrows indicate edges with lower contrast, possibly due to direct interaction of the two corresponding subunits with the substrate. All features agree qualitatively with the class in **d**, obtained from the RELION 3.0 tutorial data set. However, the features in the class from the ES-IBD sample appear smaller and more diffuse, indicating a finite degree of structural heterogeneity.

a b

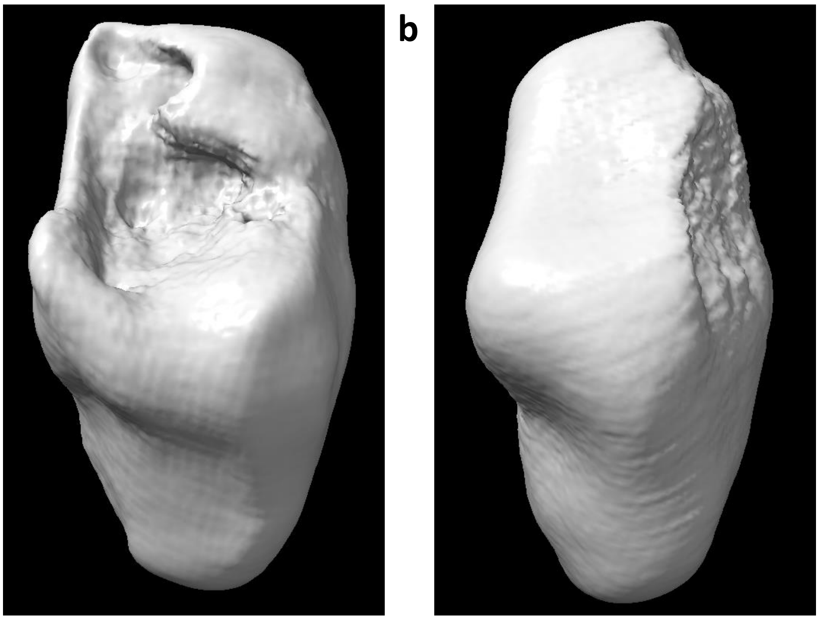

Figure S3: 3D EM density maps for ice-free ES-IBD $\beta$-gal sample obtained using C1 symmetry in RELION's automated refinement. **a** Map obtained using a subset of 50,000 particles showing localized deformation, possibly due to orientation dependent deformation on landing. **b** Map obtained from a subset of 16,400 particles, after multiple 2D and 3D classification steps, showing significantly less deformation.

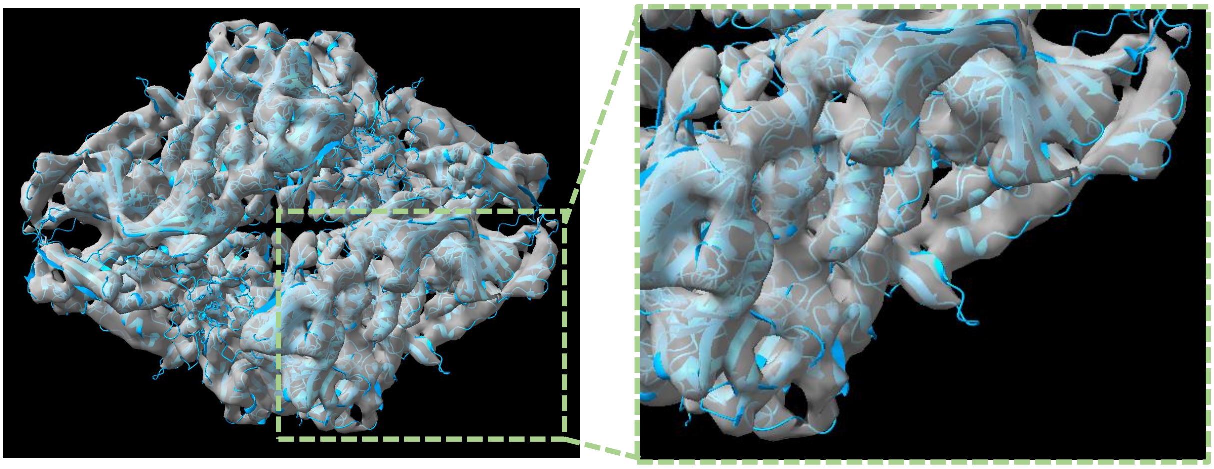

Figure S4: 3D structure from $\beta$-gal control sample in ice, obtained using the same solution as for the native ES-IBD samples (200mM ammonium acetate pH 6.9). The structure converged at a Gold-standard resolution (0.143 FSC) of 9 Å. The superimposed PDB model indicates very good agreement of secondary structure, allowing us to exclude protein preparation for mass spectrometry as a major bottleneck of the native ES-IBD workflow.

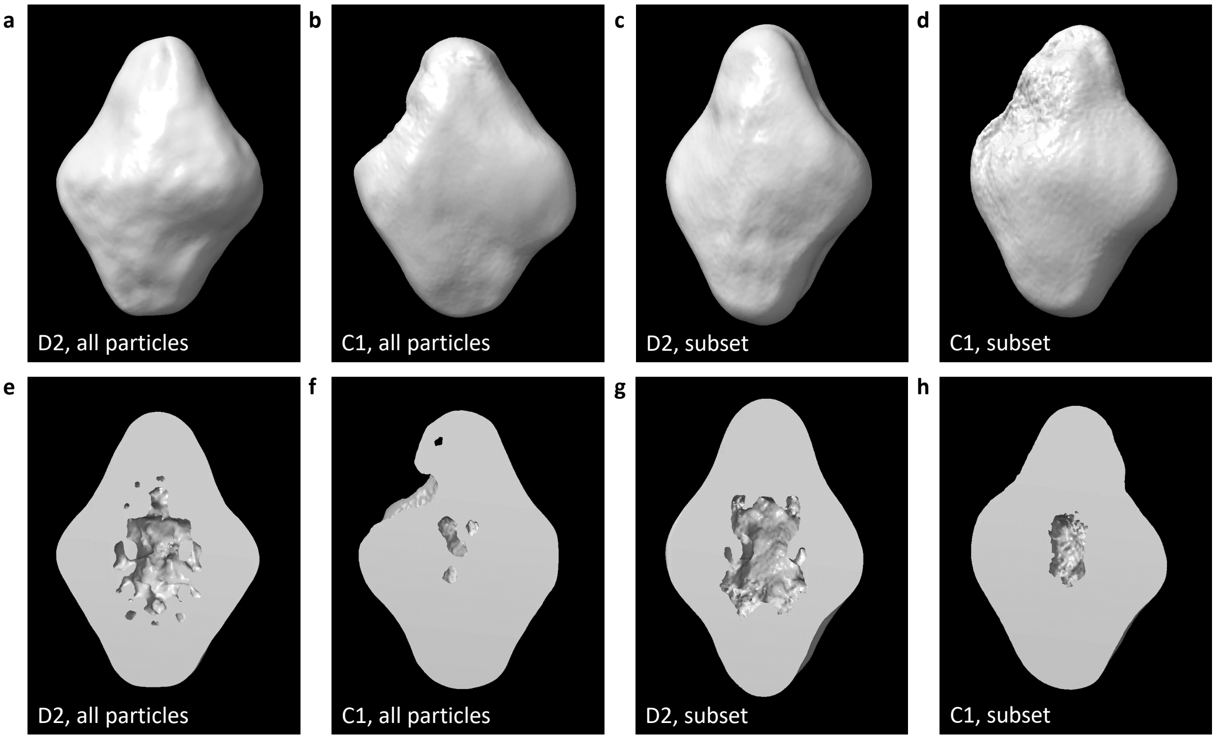


Figure S5: 3D EM density maps of the native ES-IBD sample of $\beta$-gal. The panels indicate if no symmetry (C1) or dihedral symmetry (D2) was applied during the 3D auto-refine step in RELION. Panels **a** and **b** show structures that were obtained using all 50,000 particles while panels **c** and **d** were generated using a subset of 16,400 particles. The panels in the second row show cross-sections of the structures above. See movie 1 for rotating structures.

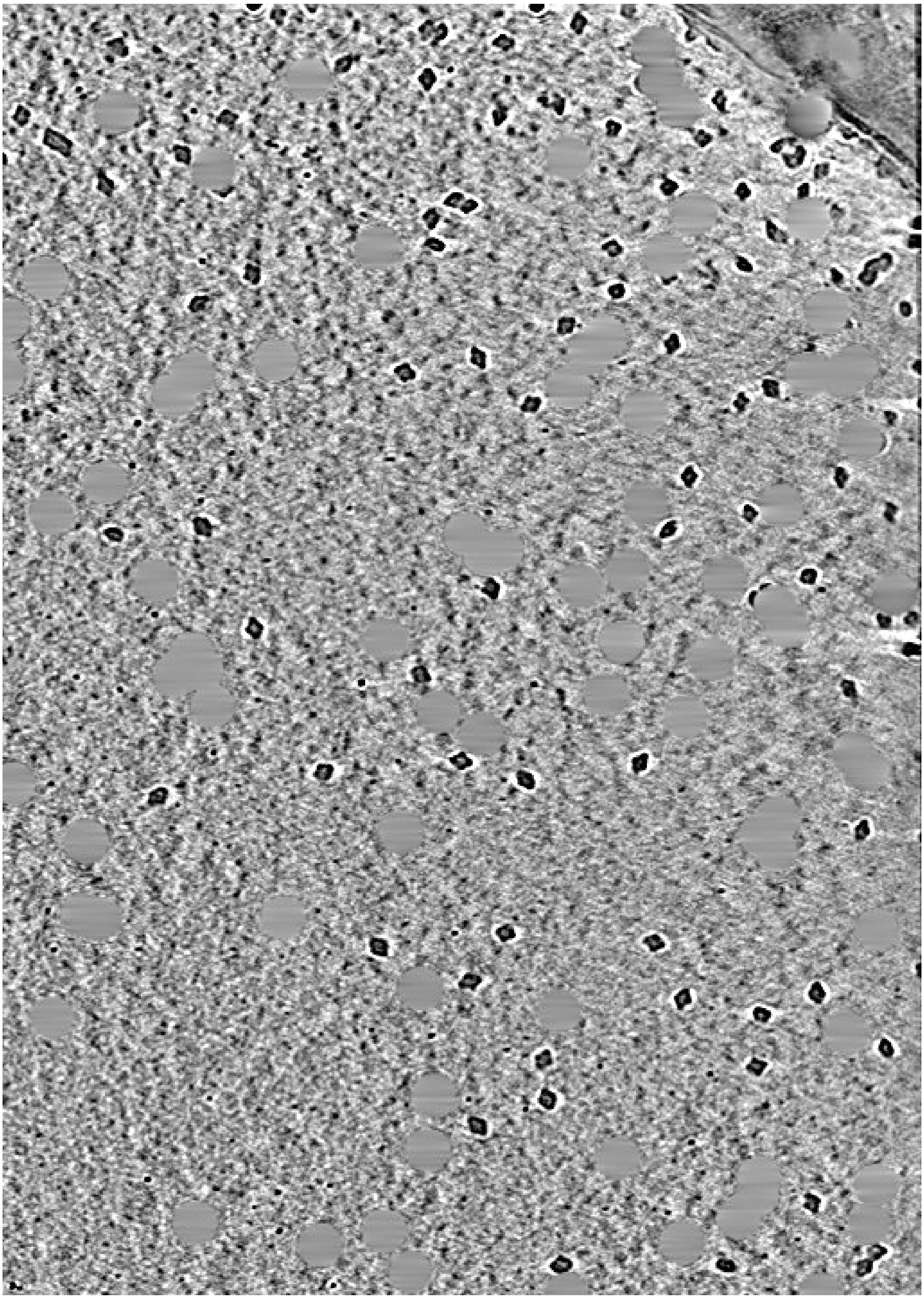

Figure S6: Tomograpic slice of a native ES-IBD sample of $\beta$-gal on a 3 nm amorphous carbon film. Shape and density distribution agree well with 2D classes and 3D maps from single particle analysis. Gold nanoparticle densities were deleted (grey circles). See movie 2 for other slices.

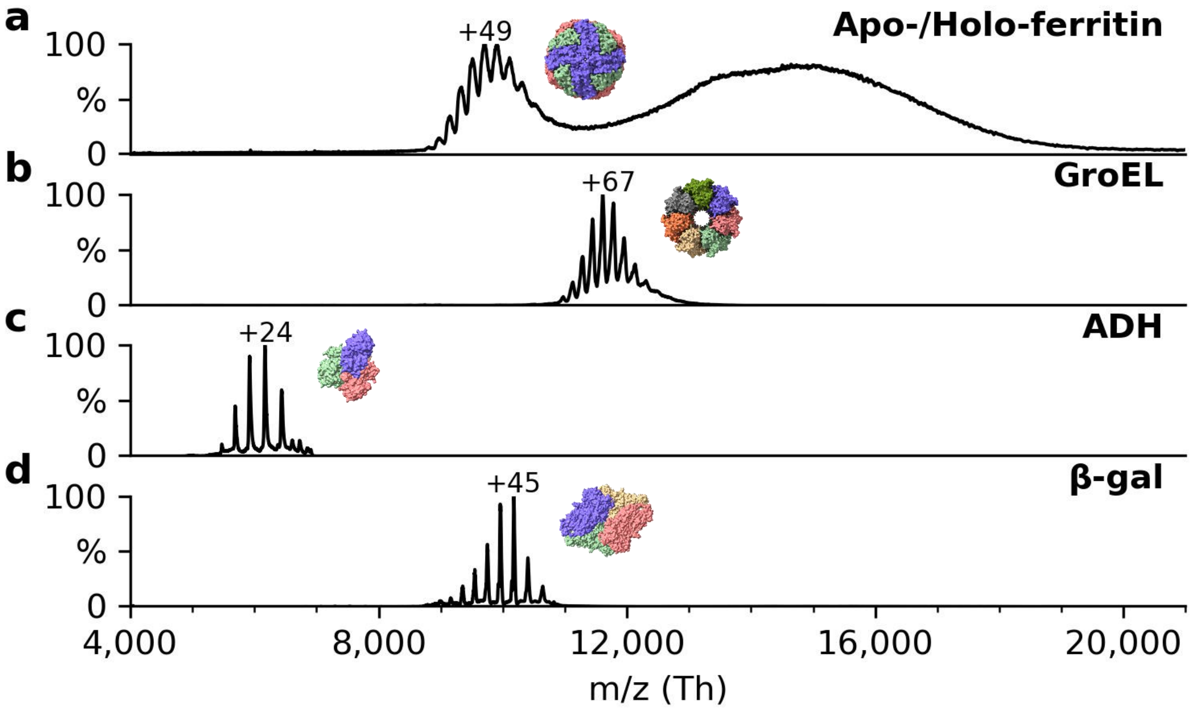


Figure S7: Non-activating, native mass spectra of apo/holo-ferritin, GroEL, ADH, and $\beta-gal$. For native ES-IBD experiments, native oligomers were mass selected, i.e., tetramer for ADH and $\beta$-gal and tetradecamer for GroEL. Both apo- and holo-ferritin were selected and deposited at the same time. The mass spectrum of holoferritin (right) shows no resolved charge states, due to the continuous size distribution of iron cores in our sample. For all other proteins, the most abundant charge states are labeled. They are much lower than typical charge states from denatured samples indicating near-native gas-phase structures.
